# Collective animal navigation and migratory culture: from theoretical models to empirical evidence

**DOI:** 10.1101/230219

**Authors:** Andrew M. Berdahl, Albert B. Kao, Andrea Flack, Peter A. H Westley, Edward A. Codling, Iain D. Couzin, Anthony I. Dell, Dora Biro

## Abstract

Animals often travel in groups, and their navigational decisions can be influenced by social interactions. Both theory and empirical observations suggest that such collective navigation can result in individuals improving their ability to find their way and could be one of the key benefits of sociality for these species. Here we provide an overview of the potential mechanisms underlying collective navigation and review the known, and supposed, empirical evidence for such behaviour, and highlight interesting directions for future research. We further explore how both social and collective learning during group navigation could lead to the accumulation of knowledge at the population level, resulting in the emergence of migratory culture.

## Introduction

Animal movement is a fundamental driver of ecological and evolutionary processes. Movement, and specifically migrations, couple disparate populations and ecosystems by transporting individuals, nutrients, pathogens and genes (***Altizer et al., 2011***; ***Bauer and Hoye, 2014***). For individuals, migrations facilitate access to spatially and temporally varying resources; however, there are significant costs and challenges associated with migration (***Dingle and Drake, 2007***). Perhaps the most serious challenge is navigation – animals must find their way through often complex environments along migration routes that can span tens of thousand of kilometers and take many months (sometimes generations) to traverse. To successfully complete these migrations, animals employ a diverse range of sensory modalities and can respond to an impressive array of cues, including magnetic fields, light polarization, landmarks, odors, and celestial bodies (***Gould and Gould, 2012***). While in some contexts the preferred navigation route is genetically encoded and instinctive, for others this must be discovered or learned from others.

Although the mechanisms of animal navigation have fascinated researchers for decades, focus has primarily been at the level of the individual (***Gould and Gould, 2012***). However, many migratory species are known to move in large groups (***Milner-Gulland et al., 2011***) and social interactions can alter migratory movement decisions (***Dalziel et al., 2016***; ***Torney et al., 2018b***). How individual navigation ability is affected by social interactions, and what unique navigational abilities can emerge at the collective level, has been far less studied, although a growing body of theoretical and empirical results supports the hypothesis that social interactions during collective navigation can lead to improved navigational ability (Fig. 1). We define *collective navigation* as the outcome of navigating within a social context. These outcomes can be beneficial, neutral, or detrimental, although we note that for the most part we, and the field in general, focus particularly on positive outcomes.

**Figure 1.**
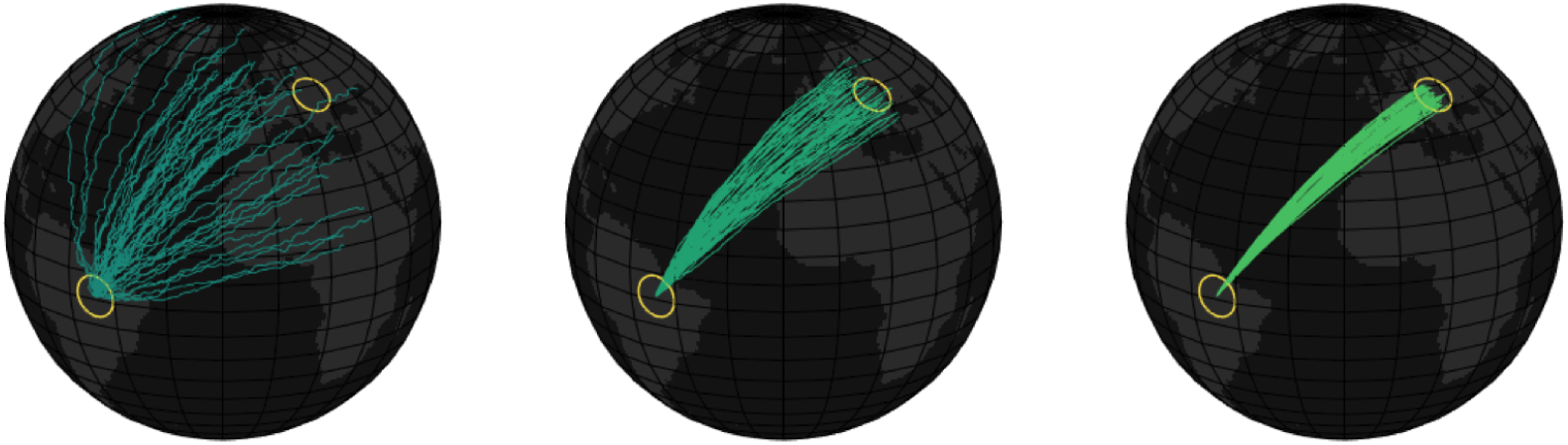
Illustration of the potential benefit of collective navigation. In this hypothetical example, migrants seek to travel from South America to Europe, with each line denoting a particular group of migrants. On average the navigation accuracy improves from left to right, which could be due to an increase in the size of the group, increase in the fraction of leaders in the group, or learning by individuals. See Box 1 for details of collective navigation mechanisms. In reality, the ‘best’ route may not be the straightest path, as navigational efficiency will be a function of several considerations, including resource distribution, perceived safety and cumulative hydro/aerodynamic efficiency.

Here we review the growing literature on collective navigation in order to: i) provide an overview of the theoretical mechanisms by which social interactions can facilitate navigational benefits; ii) synthesize empirical support for these mechanisms across several taxa, both in controlled experiments and in observations from the field; iii) explore how social and collective learning may allow for the accumulation of information at the population level, thus leading to the emergence of animal culture in a migratory context; and iv) highlight potentially fruitful directions to further the study of collective animal navigation, especially with the use of new technologies.

We describe five broad mechanisms for collective navigation: many wrongs, emergent sensing, leadership, social learning and collective learning (Box 1). The first three describe different ways in which social interactions may lead to improved navigation during a single navigational bout. Social and collective learning (see Box 1 D&E for distinction between these two) describe how information can propagate through a population or across generations, and how new information can emerge through social interactions. While previous reviews tend to focus on individual mechanisms (*e.g.*, many wrongs (***Simons, 2004***), leadership (***Conradt and Roper, 2005***) social learning (***Galef and Laland, 2005***)), here we focus on these mechanisms in the context of navigation, and highlight differences between, and interactions across, the various mechanisms. Hence, we show that the five mechanisms are not mutually exclusive, and collective navigation can be the result of a complex and dynamic set of processes spanning multiple spatial and temporal scales.

These mechanisms may also apply to many other navigational tasks in addition to migrations, across a wide range of spatial and temporal scales. For example, many animals navigate in order to discover new food sources, move up and down the water column, or locate new shelters. Because animals use environmental information to reach a specific target in these and other tasks, collective navigation mechanisms could play a role for group-living animals in improving their performance. Furthermore, while the majority of the direct empirical evidence of the collective navigation mechanisms uses birds or fish as study organisms, there are many other taxa, including ungulates, cetaceans, and insects, which navigate through their environment while traveling in groups. Where relevant, we allude to some of these less well-studied taxa as potential directions for future research.

## Theoretical models and mechanisms

The idea that the effectiveness of a collective decision-making process covaries with group size dates back several centuries, initially focusing on decision-making in humans. One classic example, from the late 18th century, is Condorcet’s jury theorem, which posited that when individuals must choose between two discrete options (e.g., the guilt or innocence of a defendant), and each jurist has a greater than 50% chance of choosing the correct option, then the accuracy of decisions will tend to improve as the size of the group increases (***Condorcet, 1976***). Later work, including***Galton(1907)***, extended this idea from discrete to continuous estimates, suggesting that the average of many independent estimates will tend to approach the ‘true’ value with increasing accuracy as group size increases – a phenomenon now known as the ‘wisdom of crowds’.

It was only much later that these ideas were adapted to non-human animal groups when, in the 1960s, researchers studying birds (***Bergman and Donner, 1964***; ***Hamilton, 1967***;***Wallraff, 1978***) and fish (***Larkin and Walton, 1969***) independently suggested that these animals could improve their navigational performance by grouping. For example, ***Larkin and Walton (1969)*** supposed that each fish within a school makes an independent estimate of the best migratory direction, and by traveling together they would tend to move in the average preferred direction of all individuals. In such a scenario, the authors found that navigation error should decrease as the inverse of the square root of the number of animals in the group, analogous to how the standard error shrinks as the sample size increases in statistical analyses due to the law of large numbers. Similarly, Condorcet’s theorem could apply in animal groups when animals must make binary or other discrete choices, such as fish ascending a river network (***Berdahl et al., 2016b***) or bees selecting a new nest site (***Seeley and Buhrman, 1999***), such that decision accuracy improves with group size in these scenarios (***Kao and Couzin, 2014***). Now known as the ‘many wrongs principle,’ the general idea that social interactions dampen individual errors is thought to be a major outcome of collective navigation (***Simons (2004)***; Box 1A).

While these relatively simple mathematical arguments provide an intuitive conceptual basis for how individuals in groups could improve their navigational accuracy, they largely ignore the complexity of the behavior of real organisms. In most animal groups, there is no entity to collate ‘opinions’ and explicitly compute the average of all individual estimates, as each individual can observe only near neighbours. Furthermore, individuals may not be equally informed about the best direction of travel, there may be complex interactions between genetically-determined and learned preferences, or group-wide biases in estimates. Because of this, it is not obvious whether navigation accuracy in animal groups would scale as these simple models predict, or whether there are limits to the real-world ability of organisms to benefit from collective navigation. More mechanistic models are necessary to shed greater light on the mechanisms underlying collective navigation in animals.

Agent-based models, where the motion of each individual is modeled explicitly in space and time (***Aoki, 1982***; ***Reynolds, 1987***) were developed in order to bridge the gap between abstract mathematical models and the behaviour of real animal groups. These models can describe how the motion of an individual is determined by its own navigational preferences, physical abilities, sensory information, and response to near neighbors. The social interaction rules are often governed by ‘zones’ of interactions, such that the response to a neighbor depends on the distance between the neighbor and the focal individual (***Vicsek et al., 1995***; ***Couzin et al., 2002***, ***2005***; ***Hein et al., 2015***). More recently, empirical data have driven the development of alternative models, where, for example, individuals respond to a fixed number of near neighbors irrespective of their distance (***Ballerini et al., 2008***), where social influence decays continuously as a function of distance (***Torney et al., 2018b***) or where interactions are modulated by considerations of the animals’ sensory capacities and limitations (***Strandburg-Peshkin et al., 2013***; ***Rosenthal et al., 2015***). Agent-based models are particularly useful because ‘experiments’ can be performed *in silico* even when the underlying equations are not mathematically tractable. Furthermore, experiments can be performed digitally to address questions that may be difficult or impossible to do with real animals in the lab or the field. For example, different parameters of the model (such as sensing ability, social interaction network, or the structure of noise) can be varied systematically, and their effect on collective navigation measured. In addition, such models allow an exploration of how collective behavior may change over evolutionary timescales (***Guttal and Couzin, 2010***; ***Hein et al., 2015***). The results of such virtual experiments can serve as testable predictions regarding which behavioral parameters are likely to be important for real animals, which can lead to more targeted experiments.

The simplest agent-based models of collective navigation assume that all individuals in the group are identical – they follow the same interaction rules and have the same level of navigational information or error, thus approximating the conditions that the many-wrongs principle typically assumes. Such simulations have demonstrated that many-wrongs averaging can readily arise from local social interactions if individuals balance their own preference with the direction of motion of their neighbors (***Grünbaum, 1998***; ***Codling et al., 2007***). Specifically, collective navigational performance is maximized when personal preference is given a low weight (***Codling and Bode, 2014***), if individuals exhibit some inertia in their movements (which serves to average an individual’s noisy compass estimates over time) (***Codling and Bode, 2016***), or if the underlying social structure is evenly distributed, rather than dominated by a few individuals (***Bode et al., 2012b***; ***Flack et al., 2015***).

For many other contexts, the distribution of directional preferences may be multimodal rather than unimodal. For example, different individuals in a group may have different preferred routes to the same location, and at small spatial scales, individuals can exhibit distinct preferred headings. In other cases, individuals may prefer altogether separate locations, such as when individuals in a breeding population choose from multiple overwintering grounds (*i.e.,* weak migratory connectivity (***Webster et al., 2002***)). In such cases, there will be a natural continuum between unimodal and multimodal distributions of preferences depending on the distance individuals are from the final location. Specifically, when locations are very far away, all individuals prefer to move roughly in the same direction (unimodal), but as the group approaches the locations preferences will begin to diverge (become multimodal). In such scenarios, simply taking the average of the preferred directions can be detrimental (there may well be no suitable habitat at the midpoint between preferred locations). Agent-based models that incorporate this diversity of preferences have demonstrated that, despite these challenges, groups are consistently able to reach consensus for one particular location. One robust result of both models and empirical data is that animal groups average when the discrepancy between preferred headings is small, but when the discrepancy is sufficiently large, the group spontaneously selects one of the possible headings (***Couzin et al., 2005***; ***Biro et al., 2006***; ***Strandburg-Peshkin et al., 2015***), typically the one preferred by the greatest number of individuals (***Couzin et al., 2005***; ***Biro et al., 2006***; ***Strandburg-Peshkin et al., 2015***) or the most strongly opinionated individuals (***Conradt et al., 2009***; ***Freeman et al., 2011***).

Another realistic extension of these agent-based models is to include two classes of individuals, informed and naïve, where successful navigation requires leadership by the informed class (Box 1B). In real animal groups, this can occur when the desired navigation direction is not genetically encoded and must be learned: the naïve individuals may be juveniles that lack experience of the route, or members of fission-fusion groups that are less knowledgeable about the local geography or other informative cues. One question that arises from these mixed groups is whether, and how, relevant information about which way to go can successfully percolate from a minority of leaders to the entire group. Effective leadership would not be explained by many wrongs, which would predict poor navigational ability in such scenarios, since it describes the averaging of estimates across the entire group. This challenge is compounded if information about who is informed cannot be directly signaled, and leadership must arise despite this anonymity. Models in which a group is composed of an informed subclass and an uninformed subclass show that surprisingly few informed individuals are necessary to effectively lead a group (***Romey, 1996***; ***Huse et al., 2002***; ***Couzin et al., 2005***), with a relatively sharp transition from ineffective to effective leadership. Models suggest that leadership can be enhanced if the informed subclass moves more quickly than the naïve majority (***Janson et al., 2005***) in order to increase their contact rate or to signal information, although this is not a requirement for effective leadership (***Couzin et al., 2005***). Further studies have shown that naïve individuals can even improve collective navigation because they contribute error that can actually stabilize consensus decision-making and increases the speed and sensitivity of consensus (***Couzin et al., 2011; Hartnett et al., 2016***).

Knowledge heterogeneity may be an outcome of evolution, rather than simply a consequence of age structure or mixing. Evolutionary simulations, in which gathering information is costly (as it necessitates, for example, developing enhanced sensory capabilities or diverting more attention to information gathering) suggest that frequency-dependent selection drives the evolution of leaders (those who predominantly rely on environmental cues) and followers (those who predominantly rely on social cues) (***Guttal and Couzin, 2010***; ***Torney et al., 2010***). This may even occur when individuals are very sparsely distributed in space, and thus rarely interact, demonstrating that individuals can benefit from "collective" navigation even if they do not appear to be grouping at all (***Guttal and Couzin, 2010***).

Differential levels of knowledge also provide opportunities for naïve individuals to learn migratory routes and other relevant information socially for use in future journeys. Such unidirectional copying behaviour is typically referred to as social learning (***Rendell et al., 2011***) (Box 1D).***Hamilton (1962)*** and others proposed the intuitive idea that young migrants could learn migration routes when traveling with more experienced individuals by being exposed to cues associated with that route. Social learning may also occur between individuals of the same age class. For example, in fission-fusion populations, there may be local heterogeneity in knowledge about the environment due to the mixing of individuals among groups. In such scenarios, animals can gain information about relevant geographical features or landmarks by following better informed, transient, group members. While the role of social learning in collective navigation has received substantial empirical support (which we discuss in a later section), there are fewer theoretical models. However, the models that do consider the transmission of information across generations suggest that it could lead to collective memory in a population, allowing for migration routes and destinations to be culturally established and maintained (***Huse et al., 2002***; ***Fagan et al., 2012***; ***De Luca et al., 2014***).

In addition to social learning, whereby information is passed from one individual to another (or several others), social interactions can also lead to collective learning, where new information emerges *de novo* as a result of social interactions (Box 1E). For example, a group can jointly discover an improved route, through many wrongs or randomly by noise injected from social interactions, which can then be learned by the group members. ***Kao et al. (2014)*** demonstrated theoretically that the collective context within which decisions are made can substantially alter what individuals learn about their environment, enabling them to maximize collective accuracy without the need for special social cognitive abilities. By altering how individuals experience the world, social interactions can affect what aspects of the environment are learned and can contribute to new knowledge within the group that improves navigation. Such learning can lead to the accumulation of increasingly better navigational solutions over time, in a process analogous to cumulative cultural evolution ***(Biro et al., 2016)***. We return to both social and collective learning in a later section, and provide more explicit suggestions for the key aspects that differentiate them, as well as for the consequences that these differences have for what form migratory cultures take.

While the above models largely presumed a preferred absolute travel direction or target, in many contexts animals navigate by following local cues. Additionally, animals may perform local search to find winds or currents that are favourable for their migration route ***(Chapman et al., 2010)***. In these scenarios, successful navigation can require detecting and climbing environmental gradients, such light, odor, temperature or current (***Gould and Gould, 2012***). In theory, a group could act as a spatially distributed sensory array spanning weak environmental gradients and amplifying weak signals (***Couzin, 2007***; ***Torney et al., 2009***, ***2011***; ***Berdahl et al., 2013***). In such a scenario, the many-wrongs effect (Box 1A) could help a group climb a noisy environmental gradient if each individual makes an independent assessment of the direction of the gradient (***Grünbaum, 1998; Codling et al., 2007***).

However, effective climbing of gradients can also occur collectively even when individuals themselves are unable to detect gradients. Known as emergent sensing, social interactions facilitate comparisons across scalar measurements made by individuals, leading to a collective computation of the environmental gradient (***Torney et al., 2009***, ***2011***; ***Berdahl et al., 2013***) (Box 1C). For example, by altering individual-level behaviour (e.g., social interactions (***Torney et al., 2009***) or swim speed (***Berdahl et al., 2013***)) in response to local *scalar* values of the environment, movement up a gradient can emerge at the group level. ***Hein et al. (2015)*** used simulations to demonstrate such group-level traits are an evolutionarily stable outcome, readily arising from selection operating on the behavior of selfish individual agents rather than explicitly on group-level properties. In contrast to the many-wrongs effect, which has a known upper-bound to accuracy, the limits of emergent group sensing are not well understood. The space of such context-dependent behavioural rules is potentially very large and much remains to be explored, both theoretically and empirically. Because current techniques to infer social interaction rules from data typically average over time and individuals they potentially miss such context-dependent behaviours that may be highly relevant to navigation.

#### Box 1. Mechanisms leading to improved accuracy during collective navigation

##### A) Many wrongs

Many wrongs is the mechanism by which a group of animals, each with a noisy estimate of the ‘correct’ navigation direction, can improve their accuracy by pooling individual estimates. At its core, it is deeply related to the law of large numbers. As long as the errors of individual estimates are not perfectly correlated with each other, and are distributed in an unbiased manner around the true value, then a simple averaging across estimates can increasingly dampen noise and home on the true value (Box Fig. 2A). Known social interactions rules have been shown to effectively average across preferences. This mechanism can operate on either continuous (such as direction of motion) or discrete (such as distinct paths or river branches) variables. In the latter case, majority (or plurality) rule serves an analogous function to simple averaging. For a group composed of individuals with differing accuracies, many wrongs may still improve accuracy, although accuracy would be maximized by a weighted average.

##### B) Leadership

Leadership results when informed individuals, which may form a small minority of the group, successfully guide naïve individuals towards favorable environments. Smaller groups may allow for individuals to recognize leaders and preferentially follow them, while in large groups, leaders are likely to be anonymous. Nonetheless, social influence can lead to successful leadership, with a surprisingly small number of leaders necessary for accurate navigation (Box Fig. 2B). Naïve individuals can even help ensure democratic decision-making, potentially aiding in a many wrongs improvement of accuracy. Who is a leader can depend on the specific context, so that over the course of a migration, leadership may be distributed among many members of the group.

##### C) Emergent sensing

Emergent sensing occurs when a group can navigate collectively even when no individual has the ability to assess the correct direction of motion. If an individual, for example, can make only scalar measurements of an environmental cue and has no memory, then it has no knowledge of the gradient of the cue. But a group can, collectively, measure and follow a gradient if the measurements made by multiple individuals can be compared. The group would then function as a distributed sensor network. Although many animals that navigate together cannot directly communicate and compare measurements with each other, context-dependent behavior (where some aspect of behavior is tied to the value of the measurement) can effectively facilitate such comparisons, even if no individual is aware of such comparisons (Box Fig. 2C).

##### D) Social learning

Social learning allows knowledge possessed by leaders to percolate through the group and across generations. If naïve individuals are led along a particular path by more knowledgeable group members, those individuals may learn about cues associated with that path, therefore becoming part of the informed subset themselves over time. Similarly, individuals with similar ages, or levels of experience, may have differing knowledge of specific routes or cues and this information may be homogenized via learning during group travel. In both contexts, the learning is unidirectional – individuals gain personal information by following others who already have that information. For navigational tasks where there is no genetically encoded preferred direction, social learning can be the primary mechanism by which navigational information persists over generations. Innovations to routes (e.g., novel shortcuts, detours) originate with leaders/demonstrators at the individual level, and can be passed on to followers/observers.

##### E) Collective learning

Collective learning is the emergence and retention of new knowledge resulting from the dynamics of social interactions. It differs from social learning in that route innovations are generated from the interaction of multiple individuals. For example, a group can improve the route that it takes through the many wrongs mechanism, and this new route can then be learned by individuals in the group. Alternatively, naïve individuals may inject random noise (stochastic factors such as sensory, or movement, errors) into a traveled route, and improved routes could be haphazardly discovered and subsequently learned - although this may require the group to also have the capacity to filter out "bad" innovations. Both collective social learning may lead to gradual improvements, or ‘ratcheting’, of the efficiency of the learned route over time.

**Figure 2.**
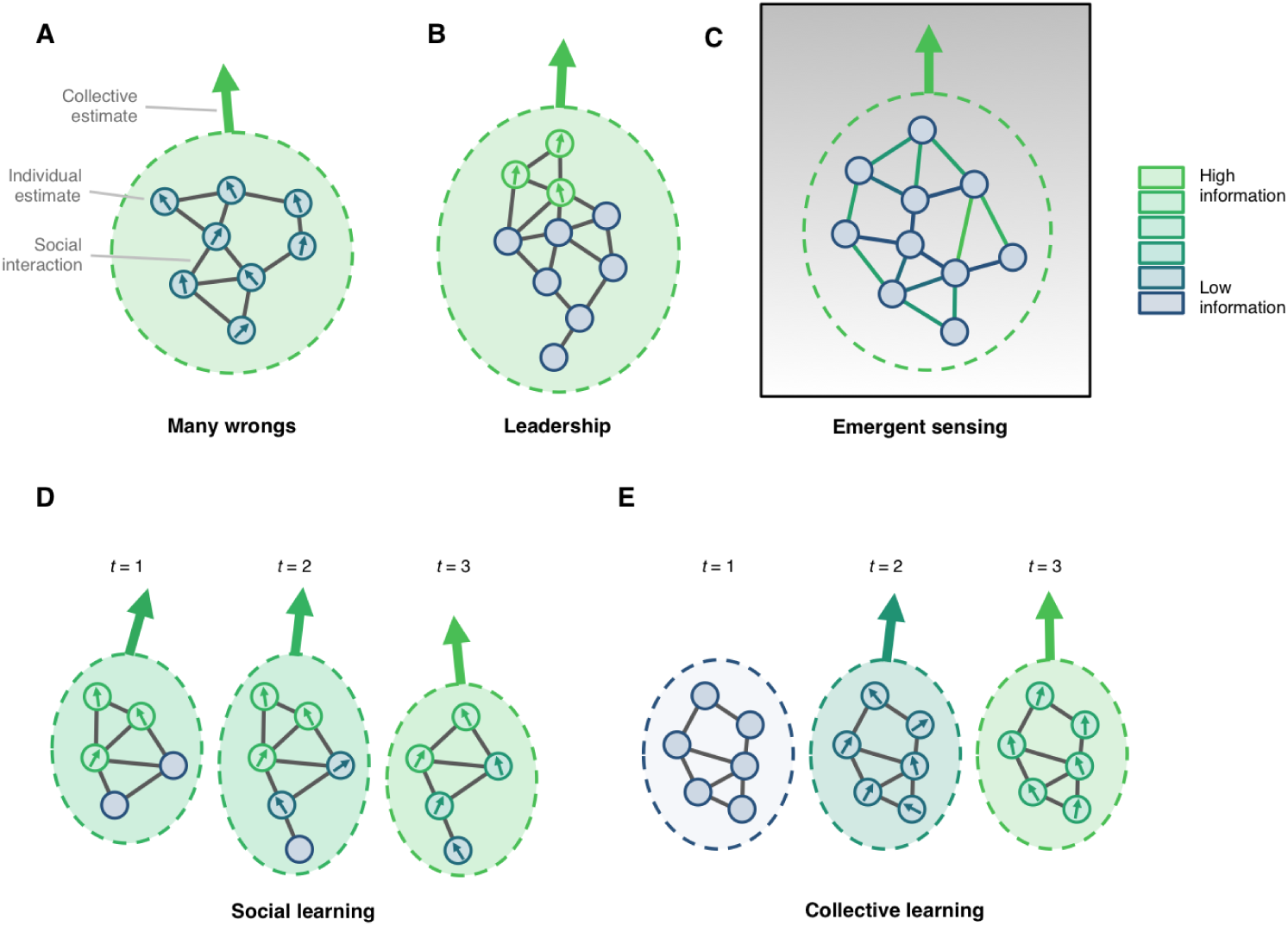
Mechanisms leading to improved accuracy during collective navigation. **(A)** Many wrongs: noisy estimates from many individuals are averaged to produce a more accurate collective estimate. **(B)** Leadership: a subset of informed individuals guides naive individuals. **(C)** Emergent sensing: comparisons of individual measurements of the environment via social interactions allows a group to detect gradients. Here information is present in the interactions (links) rather than the individuals themselves. **(D)** Social learning: navigational information passes from informed individuals to naive individuals over time. **(E)** Collective learning: new information is generated through collective processes over time. (Please note: This figure would go in Box 1.)

## Signatures in the wild

The theoretical and modeling work on collective navigation make a number of broad predictions about the movement of animals in the wild. A few prominent examples include: (i) larger groups should, on average, navigate more accurately than smaller groups; (ii) a small proportion of informed leaders should be able to effectively lead a large group; (iii) larger groups should better sense and respond to their environment and (iv) individuals should be able to learn to improve their own navigational knowledge or ability by socially facilitated exposure to relevant environmental cues. One avenue by which to study collective navigation empirically is to compare these theoretical predictions to observational data from the wild. Observations that appear to agree with these theoretical predictions would not conclusively demonstrate collective navigation in these species but would highlight potentially relevant species for further experimental study. In this section we summarize observations of real animals – primarily in migratory species – that are consistent with predicted outcomes of collective navigation (also see Table 1).

**Table 1.**
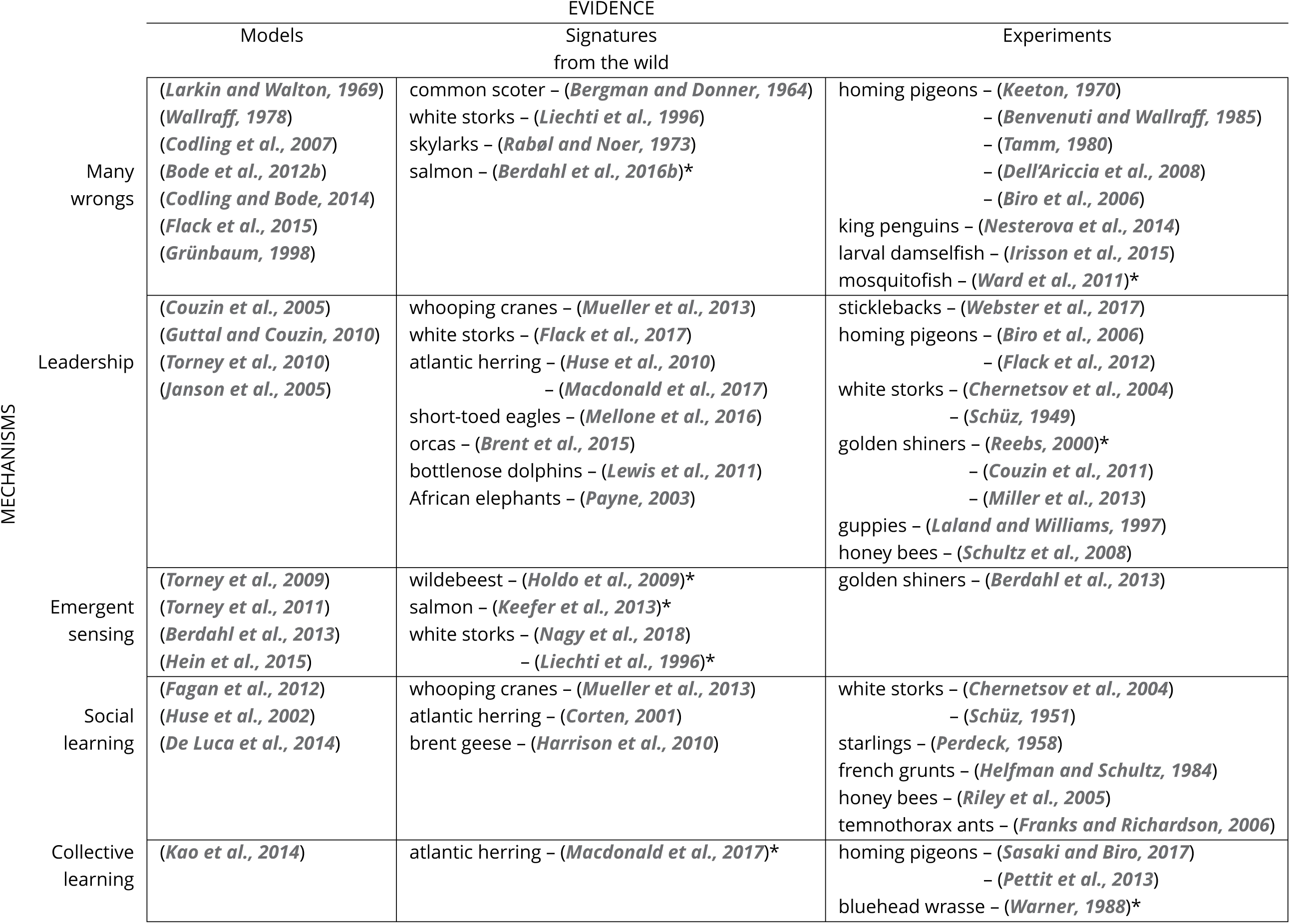
Summary of collective navigation studies categorized by the primary mechanism and type of evidence. In entries marked with an *, the exact mechanism is not clear.

The earliest observational studies focused on the many-wrongs principle (Box 1A) in migrating birds. Consistent with the predictions of this principle, directional accuracy appears to increase with group size for fowl (***Bergman and Donner, 1964***), white storks (***Liechti et al., 1996***) and skylarks (***Rabøl and Noer, 1973***), although the latter study (***Rabøl and Noer, 1973***) is limited due to a small range of group sizes. More recently, experimental studies using GPS-tracked individuals have yielded more rigorous support for many wrongs (***Dell’Ariccia et al., 2008***; ***Biro et al., 2006***) (see next section for details).

Migrations that rely on local cues for effective navigation provide support for the theory of collective environmental sensing (Box 1C). Congruent with predictions of collective sensing, storks in flocks are better than individuals at locating thermal updrafts along their migration route, which the birds use to gain altitude more efficiently (***Liechti et al., 1996***). Further, wildebeest move towards new food resources that are ostensibly beyond their personal sensory range (***Holdo et al., 2009***), although an alternate explanation is that rain clouds or lightning flashes may be visible over large distances and provide meaningful information to individuals.

We see evidence of leadership (Box 1B) in the wild, both within and between generations. The predictions of ***Guttal and Couzin (2010)*** and ***Torney et al. (2010)*** that distinct leader and follower behavioural types exist within a generation is supported by recent empirical evidence from a flock of wild white storks. ***Flack et al. (2017)*** found that during their first migration a relatively small subset of individuals act as leaders both within, and between, thermals. Leaders needed to constantly adjust their flight paths to locate regions of maximal lift within the complex physical environment of thermals, whereas followers, by exploiting social information, exhibited more efficient paths. However, these followers left thermals earlier, and at lower altitudes, resulting in them exhibiting considerably more flapping flight as they moved between thermals. In support of the idea of inter-generational leadership, ***Mueller et al. (2013)*** found that navigational accuracy increased with the age (a proxy for experience) of the oldest bird in a group, and not as a function of group size (as many-wrongs would predict) in a population of reintroduced whooping cranes (*Grus americana*). Thus, in this system, younger birds benefit from traveling with older, more experienced, birds. Similarly, experienced older and/or more dominant individuals show disproportionate leadership in group-living mammals with stable social structures, such as orcas (*Orcinus orca*) (***Brent et al., 2015***) elephants (*Loxodonta sp.*) (***Payne, 2003***) and wolves (*Canis lupus*) (***Peterson et al., 2002***). Further, ***Huse et al. (2010)*** and ***Macdonald et al. (2017)*** showed in Atlantic herring (Clupea harengus) that the establishment of new migratory destinations coincides with peaks in the ratio of first-time spawners to repeat spawners. This suggests that the large influx of naïve migrants swamps the ability of the older, informed, fish to lead – though an alternative (or additional) hypothesis is that the naïve individuals have a greater affinity to (collectively) track environmental gradients than do experienced individuals (***Macdonald et al., 2017***).

Navigating groups with inter-generational leadership can also lead to social learning (Box 1D). In fact, ***Mueller et al. (2013)***’s original generation of cranes succeeded to learn a migration route ‘socially’ from an ultralight aircraft. Although subsequent generations learning from older individuals was not directly tested, the phenomenon could be reasonably inferred from the data. Similarly, for Atlantic herring, genetic or environmental factors do not explain well this species’ annual return to specific sites to feed and breed, leaving social learning, where young individuals school with and learn from older and more experienced individuals, as the most likely explanation (***McQuinn, 1997; Corten, 2001***). Results from studies of light-bellied Brent geese (*Branta bernicla hrota*) show that most offspring chose staging and wintering sites in adulthood that were identical or very near to those of their parents, suggesting an important role of social learning of migratory routes, as limited genetic differences between migrants from different routes was observed (***Harrison et al., 2010***).

Often the specific mechanism underlying collective navigation is not apparent, but consistent patterns of generally increased navigational ability with increasing density reveal a potential signature of this process. For example, ***Keefer et al. (2013***) performed a statistical analysis of factors influencing the rate of salmon movement in various river conditions and showed that adult salmon are able to pass more quickly through artificial barriers – hydroelectric dams – at high densities. ***Berdahl et al. (2016b)*** performed a meta-analysis of the relationship between homing rates and the number (density) of migratory fish in Pacific and Atlantic salmon and found a consistent trend in which years of greater abundance of fish were associated with more accurate navigation to natal streams. These results could be the net effect of several mechanisms acting in parallel or in series: salmon may benefit from many wrongs when crossing the high seas (continuous estimates), consensus decision-making when choosing between two river tributaries (discrete options), and emergent sensing when locating the odor plume of a river estuary or entrance of a fish ladder.

An additional, albeit even less direct, line of evidence for animals benefiting from collective navigation may come from the interplay between population and migratory dynamics. Theory suggests that populations employing social navigation strategies may be prone to collapse and cease migration at low population size (***Fagan et al., 2012***; ***Berdahl et al., 2016a***). This predicted collapse is due to an Allee effect, whereby positive feedback between reduced population size and reduced benefits from collective navigation (regardless of mechanism) leads to further reductions in the population size. Indeed, sudden population collapse has been observed in many group migrating species (***Hardesty-Moore et al., 2018***). Further, migratory distance in wildebeest may be linked to population size (***Sinclair, 1995***; ***Harris et al., 2009***) and in the case of caribou, migrations have stopped altogether when population sizes became low, only to recover when the number of animals increased (***Dumond and Lee, 2013***).

## Experimental evidence of collective navigation

While field observations are typically only correlative and may be subject to a confirmation bias, controlled experiments can establish a causal link between one or more collective navigation mechanisms and the resulting performance of the group. However, even in controlled experiments it can still be often difficult to distinguish between various mechanisms (***Ioannou, 2017***). Here, we review several prominent examples of experiments that have demonstrated collective navigation, where the benefits range from transient improvements to longer lasting effects of socially-facilitated learning (also see Table 1).

The spatial scale of laboratory-based experiments is typically limited and are often only amenable to the study of smaller-scale challenges. However, many navigation tasks faced by animals operate on similar scales, and even many long distance movements are guided by a series of local interactions with the environment. Laboratory experiments can therefore shed light on the mechanisms governing collective navigation in nature. For example, ***Ward et al. (2011)*** showed that larger groups of mosquitofish (*Gambusia holbrooki*) make faster and more accurate binary decisions than do smaller groups. While the challenge in that particular experiment was to avoid predation, the general result may be applicable to migratory groups encountering binary choices, such as anadromous fish homing to a particular branch of a river network (***Berdahl et al., 2016b***). Emergent sensing can also be studied and revealed in the lab. Taking advantage of the innate preference of golden shiners to low light environments, ***Berdahl et al. (2013)*** demonstrated that the ability to climb environmental gradients increases with group size. The researchers found that when individual fish modulate their swimming speed in response to the local brightness level, taxis was induced at the group level, even though individuals had little ability to sense the gradient themselves. Lab experiments have also shown that collective navigation can emerge from the pooling of differential information across individuals. This pooling can occur for a single decision, for example if subgroups are knowledgeable about different informational dimensions (cues) and reach a consensus about an option that contains both cues (***Miller et al., 2013***), or across a series of decisions, for example from the dynamic allocation of leaders depending on which subgroup has the relevant information for that particular decision (***Webster et al., 2017***). Simple mechanisms like these may underlie a variety of, as yet poorly understood, situations in which groups navigate in response to local cues.

Experiments can also be performed outside of the lab. One fruitful method is to take advantage of the natural homing behaviour in some animals. In such cases, group size and composition can be easily manipulated and both the start and end points can be controlled, while taking place under naturalistic conditions. Early experiments using homing pigeons (*Columba livia*) showed conflicting results – some demonstrated a benefit of flocking on homing performance (***Tamm, 1980***) while others dids not (***Keeton, 1970***; ***Benvenuti and Wallraff, 1985***). However, these early studies assessed navigational performance only by examining the directional orientation of the birds at the release site (i.e., ‘vanishing bearings’) and the total time birds took to reach home, rather than the structure of complete trajectories. As such, they only provide rather crude measures of navigational performance. Such limitations have been overcome with the advent of miniature GPS technology that now provides high-resolution tracks of entire journeys, allowing for more detailed analyses of the selected routes. Using this technology, researchers have shown that pigeons in flocks tend to have straighter routes than when flying alone, suggesting that the group’s route comprises an averaged direction that is more accurate than individual estimates (***Dell’Ariccia et al., 2008; Biro et al., 2006***) - a form of many wrongs in operation. Similar homing experiments have been performed in other non-domesticated species. For example, groups of king penguin chicks (*Aptenodytes patagonicus*) returned to their crèches faster and via more efficient routes after displacement than did solo chicks (***Nesterova et al., 2014***), while larval damselfish *Chromis atripectoralis*, homing to their natal reef, swam straighter and faster in groups than they did when swimming individually (***Irisson et al., 2015***).

Homing experiments can also test whether a collective improvement can persist beyond the one-off experience of a given flock flight, by influencing individual orientational performance long-term through social or collective learning. In pigeons, naïve individuals not only follow more experienced leaders (***Flack et al., 2012***) but also socially learn the demonstrated homing routes while doing so, evidenced by their ability to recapitulate these learned routes during subsequent solo flights (***Pettit et al., 2013***). However, a single demonstration of a route seems to be insufficient for such transfer to occur (***Banks and Guilford, 2000***), with robust learning requiring repeated trips (***Pettit et al., 2013***). In addition, naïve birds have also been shown to have some influence during paired flights (***Pettit et al., 2013***; ***Sasaki and Biro, 2017***). Their presence likely injects noise into the decision-making process, which allows the group to try new routes and thus potentially discover improved navigational solutions. Such improvements can persist to subsequent flights (suggesting collective learning in operation), and may even accumulate over successive flights, even when there is continuous turnover within the group (***Sasaki and Biro, 2017***) (see also next section).

Displacement experiments during natural migrations are another useful and related technique for studying leadership as well as both collective and social learning. Typically, tagged juveniles or adults are translocated from their normal migration route or habitat, and the subsequent route or variance in route choice provides information about the navigation strategies of individuals. Early studies on both starlings (*Sturnus vulgaris*) and white storks (*Ciconia ciconia*) showed that displaced juveniles followed migratory paths that were common for conspecifics in the area where they had been displaced, indicating that displaced juveniles followed local conspecifics to their wintering grounds (***Schüz, 1951***; ***Perdeck, 1958***). Thus, the tendency to follow conspecifics tended to override the innate control of migratory path selection in both the starlings and white storks, a pattern confirmed by ***Mellone et al. (2016)*** in their study of the migration of juvenile Shorttoed Eagles (*Circaetus gallicus).* Furthermore, juvenile storks deprived of their social environment during migration, by being contained until all conspecifics have left the breeding grounds, do not migrate in their usual migratory direction but instead show much larger directional scatter (***Schüz, 1949***; ***Chernetsov et al., 2004***). These studies were repeated recently using satellite tracking technology, confirming that naïve white storks rely heavily on their social environment when selecting migratory routes (***Chernetsov et al., 2004***). The fact that no evidence is reported in these studies for established migratory routes *changing* through the presence of juveniles suggests that social learning, rather than collective learning, is the principal channel for transmission.

Leadership and social learning are firmly established mechanisms for the propagation of spatial information in eusocial insects. In honey bees (*Apis mellifera*), a surprisingly small subset (~ 5%) of informed individuals can lead an entire colony to a new nest site (***Seeley et al., 2006***). In these swarms leaders appear to exert influence by repeatedly flying through the swarm in the intended direction faster than the other bees (***Schultz et al., 2008***). Information is spread through eusocial insect colonies via various forms of social learning, often with an active ‘demonstrator’. Honey bees use so-called waggle dances to inform nest mates about the location of forage opportunities or new nest sites (***Riley et al., 2005***). Individual ants (specifically, *Temnothorax albipennis*) are even argued to ‘teach’ others about the location of suitable nest sites (***Franks and Richardson, 2006***) by leading naïve ants to relevant targets through so-called tandem runs (***Pratt et al., 2002***).

Additional evidence of leadership and social learning comes from laboratory and field studies with fish. In the lab, guppies and golden shiners follow experienced individuals to feeding sites (***Laland and Williams, 1997***; ***Reebs, 2000***; ***Couzin et al., 2011***; ***Miller et al., 2013***), with evidence in guppies that the routes persist even once the original leaders are removed (***Laland and Williams, 1997***). In displacement experiments in the field, such persistence can last for multiple years or even generations. In a classic study, ***Helfman and Schultz (1984)*** translocated French grunts (*Haemulon flavolineatum*) from their home range to an unfamiliar location in which the resident population exhibited fidelity to particular sites and took specific routes between them. The newly transplanted fish subsequently utilized the local residents’ routes and sites and furthermore continued to use them even after all residents had been removed. Since no changes to the residents’ routes were reported after the introduction of new fish, the most likely mechanism was leadership followed by social learning. Nonetheless, it is possible that over longer timescales, with the accumulation of many repeated group journeys between sites and a continuous population turnover, input from multiple individuals would combine to gradually shift routes, adding a collective learning element. Importantly, in control experiments, in which all residents were removed prior to conducting a transplant, the transplanted fish did not utilize the residents’ sites and routes, ruling out the possibility that all fish – the transplants in the previous treatment as well as the resident fish – were simply responding to the same environmental cues.

***Warner (1988)*** demonstrated similar social transmission in the choice of mating sites by bluehead wrasse (*Thalassoma bifasciatum*). When individuals from six reefs were displaced approximately 2 km away to a new reef location that had been cleared of conspecifics, they developed their own mating sites, which were shown to be a statistically random sample of suitable locations. Importantly, the observation that these new mating sites subsequently remained stable for multiple years is taken to indicate the presence of a persistent "culture" of site preference in bluehead wrasse. This brings us to our next section: the emergence of navigational culture from collective navigational phenomena.

## From collective navigation to the emergence of migratory culture

Across the examples so far discussed, the temporal scale at which individuals are influenced by others varies over many orders of magnitude. On the shortest scale, these influences may be the equivalent of ‘social information use’ (***Bonnie and Earley, 2007***; ***Rendell et al., 2011***), whereby the movement decisions of individuals – such as their direction, timing, or speed – are directly influenced by the presence and movement of fellow group members. However, these effects are transient, influencing the moment-to-moment decisions of individuals but with no longer-term consequences. This is how models of collective motion typically depict interactions - as consecutive timesteps. However, as we have discussed above, when individuals travel along a particular route, whether alone or with (and influenced by) a group, they have the opportunity to memorise cues along the route. These memories may feed back to influence navigational performance when the same task is attempted again subsequently, in effect preserving the knowledge over time, potentially over generations (***Biro et al., 2016***). Such cross-generational persistence through learning, and influenced by the animal’s social environment, meets criteria for culture: it can give rise to “group-typical behaviour patterns, shared by members of animal communities, that are to some degree reliant on socially learned and transmitted information” (***Laland and Janik, 2006***). Therefore, we now turn to the pathways through which the mechanisms of collective navigation we have discussed in this review can lead to the emergence of migratory cultures.

Figure 3 outlines our two major proposed pathways, with a potential crossover between the two providing a third. First, in systems with despotic leadership, followers have the opportunity for social learning: essentially, they are passive “observers” in the navigational task as they follow knowledgeable (or otherwise appointed) “demonstrators”. Observers memorising routes during these opportunities can lead to the transmission of navigational knowledge, and, if such transmission occurs repeatedly, migratory culture arises. This pathway is likely to operate in cases where, for example, there is little overlap between generations in terms of competence at, or knowledge of, a task, and where leadership is therefore the norm (such as first-time migrants travelling with parents). Second, when groups solve navigational tasks together, and do so through a many-wrongs or emergent-sensing mechanism, collective learning can replace social learning as the path to cultural transmission. In other words, when solutions to specific navigational problems emerge from pooling individual information-gathering or processing capacities, these collectively-derived solutions may be acquired by all of the group’s members, and to do so repeatedly over time, giving rise to culture. Third, in cases where leadership is not entirely despotic, but rather graded, input into navigational decisions from followers (albeit weighted less than input from higher-ranked leaders) may provide suitable conditions for collective (rather than purely social) learning. In sum, at the heart of all cultural phenomena are two things: (i) innovations that introduce new behaviours into a population, and (ii) mechanisms for the transmission of these behaviours. Our three pathways differ in *how the innovations arise* (i.e. through individual or collective intelligence) which in turn influences how they are transmitted (i.e. through social or collective learning, respectively).

**Figure 3.**
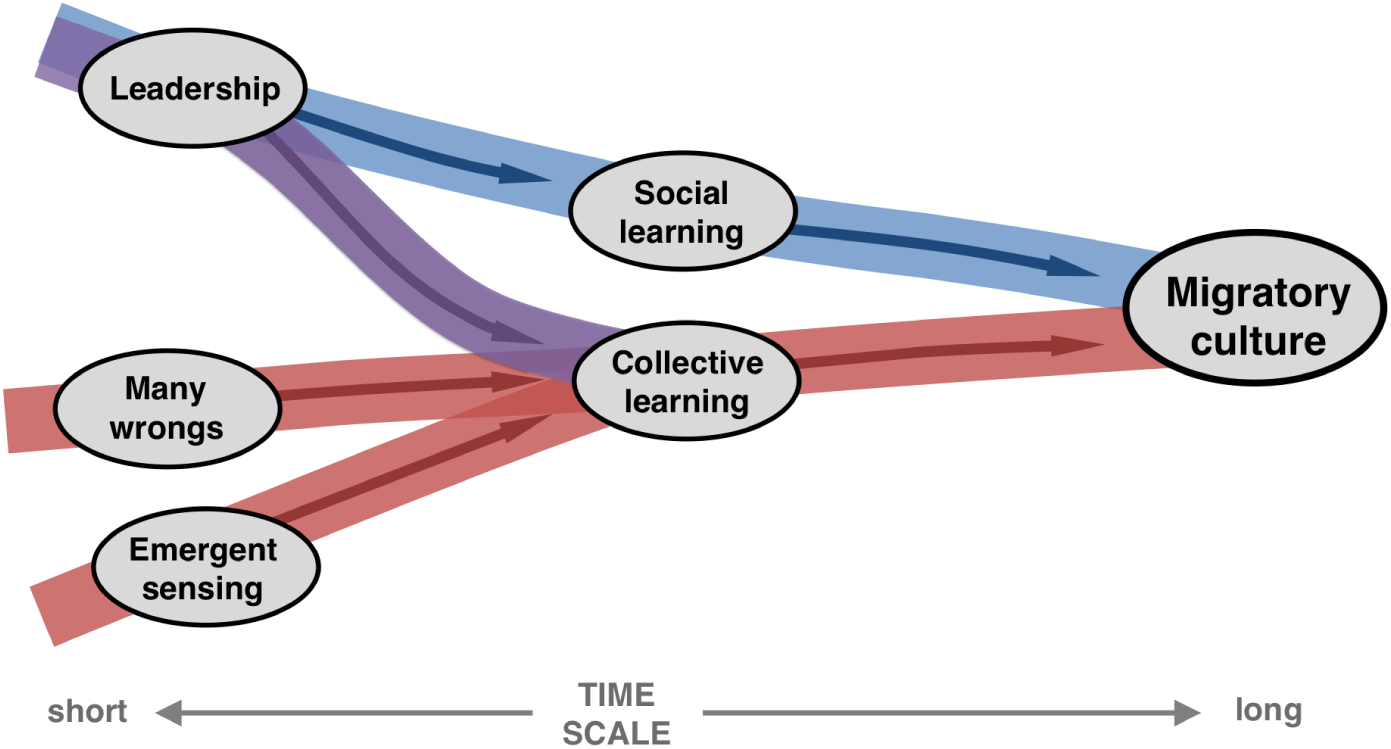
Paths to culture. Schematic summary of different pathways through which mechanisms of collective navigation may lead to navigational culture. Those mechanisms that rely on input from multiple individuals (many wrongs and emergent sensing) create opportunities for culture via collective learning, whereas social learning provides the primary pathway in groups where leadership dominates. See Fig. 2 and Box 1 for more detail on mechanisms.

Identifying examples of migratory cultures in nature is challenging. It requires multi-generational data that not only tracks the persistence of routes over time, but also confirms that they are maintained via socially mediated transmission. In other words, although it is impossible to fully discount ecological and genetic effects on route choice, these choices should demonstrably be shaped at least partially by the social environment. Furthermore, when route choice shows variation among different populations (or different co-navigating groups) of the same species, especially if moving within the same environment, this can provide important clues to cultural factors being at work. Such data are available from a small number of observational and experimental studies. In the laboratory, transmission-chain designs – a staple of experimental approaches to the study of cultural transmission (***Whiten and Mesoudi, 2008***) – have demonstrated the potential for arbitrary travel routes to be passed on via social learning along a succession of leader-follower pairs (***Laland and Williams, 1997***). In the field, natural transmission chains (such as iterative adult-juvenile joint migrations) are implicated in the maintenance of traditional travel routes (***Mueller et al., 2013***), while homing and displacement experiments mentioned previously have shown that removing older individuals or an entire resident population can cause an abrupt shift to completely different routes, and even demonstrate that a sufficient number of experienced individuals is necessary for the intra- and inter-generational stability of routes (***Huse et al., 2002; Mellone et al., 2016; Warner, 1988; Helfman and Schultz, 1984***). (In an interesting parallel, such demographic effects feature prominently in the modelling of cultural gain, drift and even loss in human technological evolution (***Henrich, 2004***)).

Individuals in groups do not need to have identical knowledge about the environment, potentially expanding the amount of information available to a group beyond the memory capacity of a single individual. With collective learning, there can be feedback between collective decisions and individual learning (individuals learn about what they experience, and what they experience is affected by the preferences and decisions of others), such that individuals in the same group may actually not learn identical representations of the same environment (***Kao et al., 2014***; ***Flack and Biro, 2013***). This can lead to a ‘collective memory,’ whereby the environment is represented in the group in a distributed manner. This distributed information can then be accessed, for example, by the dynamic allocation of leaders as different informative cues arise during navigation, as discussed earlier (***Webster et al., 2017***; ***Miller et al., 2013***).

The external environment can also help to reinforce particular routes by serving as a substrate on which memories can be encoded. For example, animals on the move can wear down the vegetation and create clear paths through the landscape. Because following these paths can be less energetically costly than generating a new one *de novo*, subsequent animals often adopt existing paths, further demarcating them. Olfactory cues left in the environment can also indicate the route taken by others (***Strandburg-Peshkin et al., 2017***). Stigmergic mechanisms such as these provide a means of social information transfer among individuals separated in time, potentially allowing for extended influence to other conspecific groups or even different species ***(Sridhar and Guttal, 2018).***

While some routes can be highly entrenched (by persisting relatively unchanged over long time scales), other paths may be further modified and improved. This gradual improvement in the efficiency or complexity of behaviour is referred to as cumulative culture ***(Boyd and Richerson, 1996)***, conceived to operate via a ‘ratchet effect’ (***Tomasello et al., 1993***) where beneficial variants are retained in the population until even more beneficial variants arise. In our schematic in Figure 3, all of our proposed pathways can lead to such increasingly better navigational solutions over repeated rounds of innovation, retention and transmission. The fact that many-wrongs and emergent-sensing are able to generate information that no individual may be capable of generating on its own (i.e. these mechanisms rely on collective intelligence) suggests that they may create either overall more effective culturally transmitted traits or may generate them faster than the pathway through individual innovation, leadership, and social learning. Nonetheless, both pathways suggest an important role for turnover in group membership in providing the ‘noise’ necessary for increasingly superior navigational solutions to emerge over time.

Cumulative culture is frequently claimed to be a human-unique trait (***Tennie et al., 2009***; ***Dean et al., 2014***), absent from other species through necessitating a suite of sophisticated socio-cognitive functions the combination of which only humans are argued to possess. To tackle the validity of this assumption, ***Sasaki and Biro (2017)*** have replicated a design previously used to study cumulative culture in humans experimentally (***Caldwell and Millen, 2008***), but with navigating pigeon flocks. The researchers removed and replaced birds in co-navigating pairs in stages, all tasked with finding a homing route from a specific release site, and found that flocks gradually improved their navigational performance across *generations,’ reaching greater efficiencies than any control individual was capable of reaching on its own. In other words, knowledge about increasingly better travel routes appeared to accumulate through collective learning, and be passed on horizontally between individuals in groups and also vertically across generations through social learning.

Thus we find signatures not only of culture, but also of cumulative culture in the development and maintenance of animal travel routes. Nonetheless, many open questions remain as to the true scope of such examples, both taxonomically and in terms of interactions with ecological and genetic effects. If present, cultural processes can have far-reaching consequences on a species’ ecology and evolution. For example, when cultural differences between groups include the emergence of distinct migratory travel routes and strong migratory connectivity between breeding and overwintering grounds (***Webster et al., 2002***), they may play a role in driving and maintaining reproductive isolation between sub-populations (***Harrison et al., 2010***), potentially affecting the evolution of the species.

Can we make predictions regarding in which species, contexts or on what scales we might expect to find migratory cultures? We suggest that a number of factors may promote the phenomenon. The ability to learn (either socially or collectively) in the context of collective movement is an essential prerequisite, as is a social structure that promotes the repeated mixing of less and more informed individuals (e.g. overlapping generations). The need to navigate to and from targets that are relatively persistent over time (e.g. to long-distance migratory destinations rather than to ephemeral food patches), but which can be reached by multiple selectively neutral alternative paths, is also likely to facilitate the emergence of stable, socially transmitted travel routes. Since local cultural innovations - points of origin for inter-group variation - can arise either from individual invention or from collective intelligence, every pathway we illustrate in Figure 3 has the potential to support cultural evolution. For migratory cultures to become cumulative, we suggest that what is important is the capacity to transmit routes with sufficiently high fidelity to enable beneficial modifications to accumulate gradually, in a ‘ratchet’-like fashion (***Tomasello et al., 1993***). Such highfidelity transmission may require (i) individual cognitive capacities to memorise landscape or other navigational cues in sufficient detail to recapitulate previously travelled routes, (ii) environments that provide such cues at sufficient resolution, and (iii) terrains that permit some degree of open-endedness in route structure.

## Outlook & future directions

Now is an exciting time for studying collective navigation. Although in this review we have emphasised empirical results, currently the theoretical predictions of collective navigation far outweigh empirical demonstrations. However, this asymmetry is already being eroded by emerging technologies, such as micro GPS tags, acoustic cameras, computer vision, UAVs and remote sensing satellites (***Hughey et al., 2018***). These technologies allow for the quantification of animal movement data at extremely fine spatial and temporal scales, and in many cases it is possible to simultaneously capture the trajectories of every animal in a group in the wild. Additionally, new technologies enable us to quantify to an astonishingly fine scale the physical environments in which these animals are moving (for example on order of ~1 cm (***Fraser et al., 2016***)). Complementing these new technologies, analytical techniques have been developed to use the data to infer the nature of social interactions (***Torney et al., 2018b***) and leadership structures (***Strandburg-Peshkin et al., 2018***) within groups, and also to explore the simultaneous effects of environmental and social drivers of collective movement (***Bode et al., 2012a; Dalziel et al., 2016; Strandburg-Peshkin et al., 2017; Calabrese et al., 2018***). In the context of collective navigation, many open questions remain (Box 2) and we are poised to make landmark discoveries – principally in understanding how animals combine social and environmental cues to find their way when navigating *through their natural habitat.*

#### Box 2. Open questions for future research

##### Do mechanisms correlate with navigational cues or life histories?

To what extent, and how, do navigational cues (e.g., magnetic field vs. landmarks) and life histories (e.g., semelparity vs. iteroparity) determine which collective navigation mechanisms animals use?

##### What are the mechanisms underlying distributed sensing in the wild?

UAVs and other new technologies allow us to capture fine-scale simultaneously trajectories of many group members (***Torney et al., 2018b; Hughey et al., 2018***) and at the same time quantify the environment in which those animals are moving with fine details (***Fraser et al., 2016; Strandburg-Peshkin et al., 2017***). Combining these technologies will allow us to explore animals combine environmental and social information when making navigating in the wild.

##### Do migratory insects benefit from collective navigation?

There are many migratory insects (***Holland et al., 2006***). Many travel at high densities and thus may benefit from collective navigation (***Hu et al., 2016***). Further, they might benefit from collective navigation even when not at high densities (***Guttal and Couzin, 2010***). With the possible exception of locusts, the role of social interactions in long-distance insect navigation is not well understood.

##### Do animals benefit from collective decision-making to optimally time their migrations?

Correctly timing a migration is vital for survival in many species (e.g., ***Satterthwaite et al. (2014)***). Just as each individual may have an independent estimate of what direction to go, each individual might have an independent assessment of *when* to go. Social interactions do influence the timing of migration behaviour (***Helm et al., 2006; Berdahl et al., 2017***). Time is distinct from space in that it is one-dimensional and asymmetric, yet many of the mechanisms for spatial collective navigation (Box 1) may have temporal analogs that could help social migrants optimally time their migrations.

##### How do collectively moving individuals sort into destination-specific groups?

To benefit from collective navigation, presumably individuals must be intending to go to the same target as the other individuals in the group, yet many fission-fusion populations mix, for example on their wintering grounds. How do animals know when to average disparate headings and when to split up? When they do split up, how do animals effectively sort into destination-specific groups?

##### What are the population genetic signatures of collective navigation?

Collective navigation on breeding migration is predicted to lead to density-dependent dispersal (***Berdahl et al., 2016b***). An exciting possibility is that resulting density-dependent dispersal may leave a population-genetic signature, which has yet to be worked out, but that could help identify the importance of social processes during navigation from genetic data alone.

##### What is the relationship between population density and group size?

The positive feedbacks between declining population size and reduced collective navigation stem from an assumption that as populations decline so will group sizes. However, it is unknown whether as population density decreases if there are fewer groups (of the same size) or a similar number of smaller groups.

##### What are the population- and ecological-level consequences of collective navigation?

Theory suggests that migratory populations reliant on collective navigation may be prone to sudden population collapse and hysteresis (***Guttal and Couzin, 2010; Fagan et al., 2012; Berdahl et al., 2016a***). Empirical tests of these predictions (e.g., ***Hardesty-Moore et al. (2018)***) could yield important insights for conservation and management.

##### How will collective navigation shape adaptation (or not) to the Anthropocene?

How will collectively navigating species fare in a world that is ever more affected by human activities, including temperature shifts, pollution and reduction and fragmentation of habitat? Will pollutants masking natural odors make collective navigation more important? Do they have the potential to alter social behaviour enough to disrupt important social processes such as collective navigation (***Brodin et al., 2013***)? Will human development lead to ‘navigational traps’ (***Sigaud et al., 2017***) Will collective navigation help or hinder species adapting to changes in optimal timing and location of migrations (***Keith and Bull, 2017***)?

##### Is there *cumulative* migratory culture in non-human animals?

We see evidence of animal migratory culture (***Helfman and Schultz, 1984; Warner, 1988; Corten, 2001***) and experiments suggest that it can even exhibit cumulative improvement in efficiency overtime (***Sasaki and Biro, 2017***), but can we find evidence for such cumulative navigational culture in natural populations? Furthermore, in line with widely used definitions of cumulative culture (e.g., ***Dean et al. (2014***)), do we also see evidence of increases in the *complexity* of the knowledge that is transmitted? Could, for example, collective memory allow migrating populations to incorporate a greater number of landmarks into a learnt route than what any one individual could memorize?

Many group-moving taxa are under-explored in terms of collective navigation. Moreover, for taxa that have been investigated the data has often been indirect or in an artificial setting. The emerging technologies described above should allow for direct exploration of the mechanism(s) underlying collective navigation in a wide range of taxa including cetaceans, marine fishes, bats, and ungulates all of which migrate and forage in large groups. Beyond increasing our understanding of their life histories, this may reveal additional mechanisms leading to group-level navigation and search. Another nearly completely unexplored taxon in terms of collective navigation is invertebrates - with the exception of the eusocial insects. Butterflies (***Brower, 1985; Frey et al., 1992***), dragonflies (***Russell et al., 1998***), locusts (***Hansen et al., 2011***) lobsters (***Herrnkind et al., 2001***), among others, travel in large groups or at high densities (***Holland et al., 2006; Hu et al., 2016***); however, to our knowledge if, and how, they might benefit from collective navigation have not been addressed. Migratory insects may benefit from many-wrongs when selecting a migratory direction, from improved collective decision-making when deciding when weather conditions (e.g., wind direction) are favourable for efficient travel, or from emergent sensing when selecting the altitude with optimal winds. We hypothesize that context-dependent social behaviour may also contribute to desert locusts’ and mormon crickets’ ability to navigate out of nutritionally depleted areas. These insects, which are normally herbivorous, turn to cannibalism when local vegetation is severely depleted (***Simpson et al., 2006***). This switch to cannibalism dramatically alters social interactions (***Buhl et al., 2006***). The allure of a nutritious abdomen in front and the threat of being bitten from behind tend to polarize these swarms into a forced march (***Bazazi et al., 2008***). Individual locusts exhibit diffusive movement, which has displacement that scales as the square root of time. In contrast, the polarized groups would travel in straighter paths (***Romanczuk et al., 2009***) – i.e. ballistic movement, which has linear displacement. Thus even if incidental, this emergent collective effect could function to move locust populations out of barren areas more rapidly, and provide another fitness benefit for cannibalism (***Hansen et al., 2011***).

As animals travel, even during goal-oriented movement such as long distance migrations, navigational accuracy will not be the only selective pressure they face. In addition to navigation, animals in nature often must simultaneously balance multiple tasks while migrating, including foraging, predator avoidance, and optimal energy allocation. Animals are effectively moving through complex topographies of risk, foraging opportunities, energy expenditure and physical terrain, and so their optimal movement will reflect some balance of all of these constraints along with their eventual intended target. Thus, the assumption that the shortest path between two points is the most beneficial may be incorrect. The ultimate goal of researchers should be to integrate navigation with natural history, ecology, aerodynamics and geography when linking fine-scale (collective) movement decisions to long-range travel (***Torney et al., 2018a; Nagy et al., 2018***).

An outstanding challenge is to link a mechanistic understanding of collective navigation to population- and ecological-level processes. Explicitly considering collective effects may dramatically change predictions of models currently used to inform management and conservation ***(Berdahl et al., 2018***). For example, sudden population collapse and hysteresis are predicted by (phenomenological) models in which migration success is dependent on social learning (***Fagan et al., 2012; De Luca et al., 2014***), leadership (***Guttal and Couzin, 2010***) and many wrongs or emergent sensing (***Berdahl et al., 2016a***). Such predictions are consistent with empirical data suggesting that population size and migratory status are linked (***Bolger et al., 2008***) and population collapse is associated with group travel in birds and fishes (***Hardesty-Moore et al., 2018***). On the other hand, collective navigation could lead to density-dependent dispersal (***Berdahl et al., 2016b***), and models predict that this density dependence should increase the robustness of metapopulations ***(Yeakel et al., 2018***). Collective navigation may also strongly affect genetic mixing within a population, by modulating the degree of migratory connectivity between breeding grounds and overwintering grounds (***Bauer et al., 2016; Turgeon et al., 2012; Brown Gladden et al., 1997; Carroll et al., 2015***), or the degree of partial migration (***Barnowe-Meyer et al., 2013; Chapman et al., 2011***). In the context of a changing climate, the cultural transmission of migration routes and destinations across generations can contribute to conservative and inflexible behavior, minimising the ability to bet-hedge in an increasingly unpredictable climate, although the social learning of adaptive innovations within a generation can also yield a greater ability to adapt to change ***(Keith and Bull, 2017***).

The study of collective behaviour typically focuses on the benefits of collective behaviour, but there may be cases where it is maladaptive. Good decision making in one context may be poor in another. Specifically, if a collective navigational strategy evolved to match a specific environment, anthropogenic modifications to that environment could disrupt the benefits of collective navigation and even make it a harmful strategy in the modern world. Indeed, ***Sigaud et al. (2017)*** revealed that, in a plains bison (*Bison bison bison*) population, information transfer mediated by fission-fusion dynamics – which presumably historically transmitted beneficial information about foraging areas – in contemporary times accelerated that population’s use of an ecological trap, triggering a precipitous population decline. Along those same lines, **Lemasson et al. (2014)** showed that schooling may impede the downstream passage of juvenile anadromous fish through artificial barriers, increasing the time they spend in this highly risky novel habitat.

Collective navigation applies not only to large-scale orientational tasks such as migrations but also to a wide range of other behavioural contexts. Navigation is important for locating new sources of food, seeking new shelters, or any other task where animals must use noisy environmental information to make decisions about where to go. Additionally, although the mechanisms may be different, there are likely rich parallels between collective search in animals and collective sensing in single-celled organisms and even groups of cells within an organism (***Haeger et al., 2015***). Finally, all of these biological systems may yield mechanisms, ‘discovered’ by eons of evolution, that could provide lessons and inspiration for human technologies (***Brambilla et al., 2013***), such as swarm robotics and particle swarm optimization.

## Author contributions

All authors participated in the review and contributed to this manuscript.

## Acknowledgments

AMB is grateful to Colin Torney for conversations that inspired and shaped this project. We also thank Thomas Mueller and two anonymous referees for helpful comments on an earlier version of the manuscript. This paper is a direct result of a workshop funded by NSF grant IOS-1545888 and the Santa Fe Institute.

